# Heat stress and recovery induce transcriptomic changes in lactogenic bovine mammary epithelial (MAC-T) cells

**DOI:** 10.1101/2024.05.15.594241

**Authors:** Xingtan Yu, Rebecca M. Harman, Nikola Danev, Guangsheng Li, Yifei Fang, Gerlinde R. Van de Walle, Jingyue Ellie Duan

**Author notes:** Corresponding author: Jingyue Ellie Duan. Authors’ email addresses: Xingtan Yu; Rebecca M. Harman; Nikola Danev; Guangsheng Li; Yifei Fang; Gerlinde R. Van de Walle; Jingyue Ellie Duan.

## Abstract

Heat stress (HS) in cattle significantly challenges the dairy industry by reducing milk production. However, the molecular mechanism behind mammary gland responses to HS and recovery remains poorly understood. This study aimed to determine the transcriptomic changes in lactogenic bovine mammary epithelial (MAC-T) cells after HS and post-HS recovery. Six culture conditions were analyzed: MAC-T cells cultured in basal medium, cells in lactogenic medium to induce differentiation, differentiated cells at standard temperature (37℃) or HS (42℃) for 1 hour. HS cells were collected after incubation at 37℃ for either 2 or 6 hours to examine the extent of recovery.

A total of 1,668 differentially expressed genes (DEGs) were identified. Differentiated cells expressed genes associated with milk lipid synthesis, indicating lactogenic potential. HS suppressed genes involved in cellular differentiation and activated heat shock protein genes. Several transcription factors were identified as potential regulators of HS response. During recovery, chaperon-mediated protein folding genes remained elevated. Apoptosis regulation genes were induced at 2 hours, and cellular homeostasis regulation genes were enriched at 6 hours. Overall, these findings provide a foundation for the molecular mechanisms involved in HS and recovery in cattle.

## Introduction

Heat stress (HS) has negative impacts on milk production and the overall well-being of dairy cattle, resulting in significant economic losses globally ^1,2^. HS activates a heat stress response (HSR), an evolutionarily conserved, intrinsic, cellular self-protective mechanism observed across mammals. Within this response, a cohort of protein chaperones, named heat shock proteins (HSPs), undergoes robust induction to maintain the structures of other proteins, and as a result, cellular homeostasis ^3^. In mouse models, the HSR triggers rapid and extensive transcriptional changes across thousands of genes ^4^.

To investigate the HSR in dairy cattle ^5,6^, high throughput RNA sequencing (RNA-seq) has revealed altered gene expressions associated with inflammation and metabolisms in peripheral white blood cells ^7^ and liver ^8^ in dairy cows under HS. Also, gene signaling pathways related to glucocorticoid biosynthesis and apoptosis were found enriched in bovine oocytes collected during summer ^9^. In mammary gland tissues of HS cows, transcriptomic ^10^ and metabolomic ^11^ approaches have identified the downregulation of casein genes, amino acid transporter genes, and lipid metabolism pathways, whereas upregulation has been observed in HSPs, immune responses, and inflammatory responses.

MAC-T cells, established from immortalized bovine primary mammary epithelial cell (MECs) ^12^, can be induced to a lactogenic state to mimic the transcriptomics of the lactating udder^13^. Previous studies using MAC-T cells under HS conditions have primarily focused on their undifferentiated state. For example, expression of HSP genes and apoptosis-related genes was significantly upregulated in undifferentiated MAC-T after HS treatment ^14^, whereas baseline expression of milk protein genes was downregulated ^15^. Recent work employing RNA-seq and methylated RNA immunoprecipitation sequencing (MeRIP) profiled transcriptomics and N6-methyladenosine (m6A) modification of HS undifferentiated MAC-T cells ^16^, revealing upregulated expression of heat shock factor 1 (*HSF1*), heat shock protein 70 (*HSP70*), and apoptosis-related genes, and altered m6A modifications within genes in TGFβ, Wnt, and Hippo signaling pathways. Currently, the impacts of HS on lactogenic MAC-T cells and milk synthesis pathways remains poorly understood, and few potent regulators or pathways have been identified^14,15^.

The residual impact of HS during the post-HS recovery phase in dairy cows has been explored to a limited extent through *in vivo* or *in vitro* models. In lactating dairy cattle, a continued decline of dry matter intake (DMI), milk yield, and milk proteins, has been observed during a five- day recovery period after moderate HS ^17^. In another study, no residual effect of HS was detected on protein production in milk after 7-day recovery ^18^. Primary bovine mammary epithelial cells exposed to HS for 1 hour demonstrated a gradual return to baseline levels of HSP genes and casein genes after a 24 hour recovery, yet the expression of heat shock protein 27 (*HSP27*) and casein alpha subunit 1 (*CSN1A1*) remained significantly higher than pre-HS levels ^19^. These intriguing results were obtained using qPCR to examine the expression of select genes, warranting global gene expression analysis, to better understand the recovery process of heat stressed mammary epithelial cells.

In this study, we aimed to profile the transcriptomic changes of lactogenic MAC-T cells subjected to HS and after recovery periods. Lactogenic MAC-T cells were exposed to HS (42°C) for 1 hour and allowed to recover at 37°C for intervals of 2 or 6 hours. Using RNA-seq, we identified differentially expressed genes and conducted functional enrichment analyses. The results from this study revealed transcriptomic dynamics during HS and recovery processes and provided us with a list of potential regulators contributing to HS response and recovery mechanisms.

## Results

### RNA-seq statistics summary

Experimental design of the present study was shown in Fig. 1. The statistics of RNA-seq data are summarized in Additional file 2: Table S2. Overall, our sequencing data was of high quality as indicated by quality values (Q20 > 98% and Q30 > 94%) and average unique mapping rates above 96%. The correlation of samples was determined by principal component analysis (PCA) using the top 300 variable genes. After batch effect removal, samples clustered according to treatment group (Fig. 2). A total of 1,668 differentially expressed genes (DEGs) were identified from six pair-wise comparisons (Table 1). The highest number of DEGs (n = 472) was found in the comparison HS-D vs. D6R and the lowest number of DEGs (n = 151) was found in the comparison HS-D vs. D2R.

**Figure 1.**
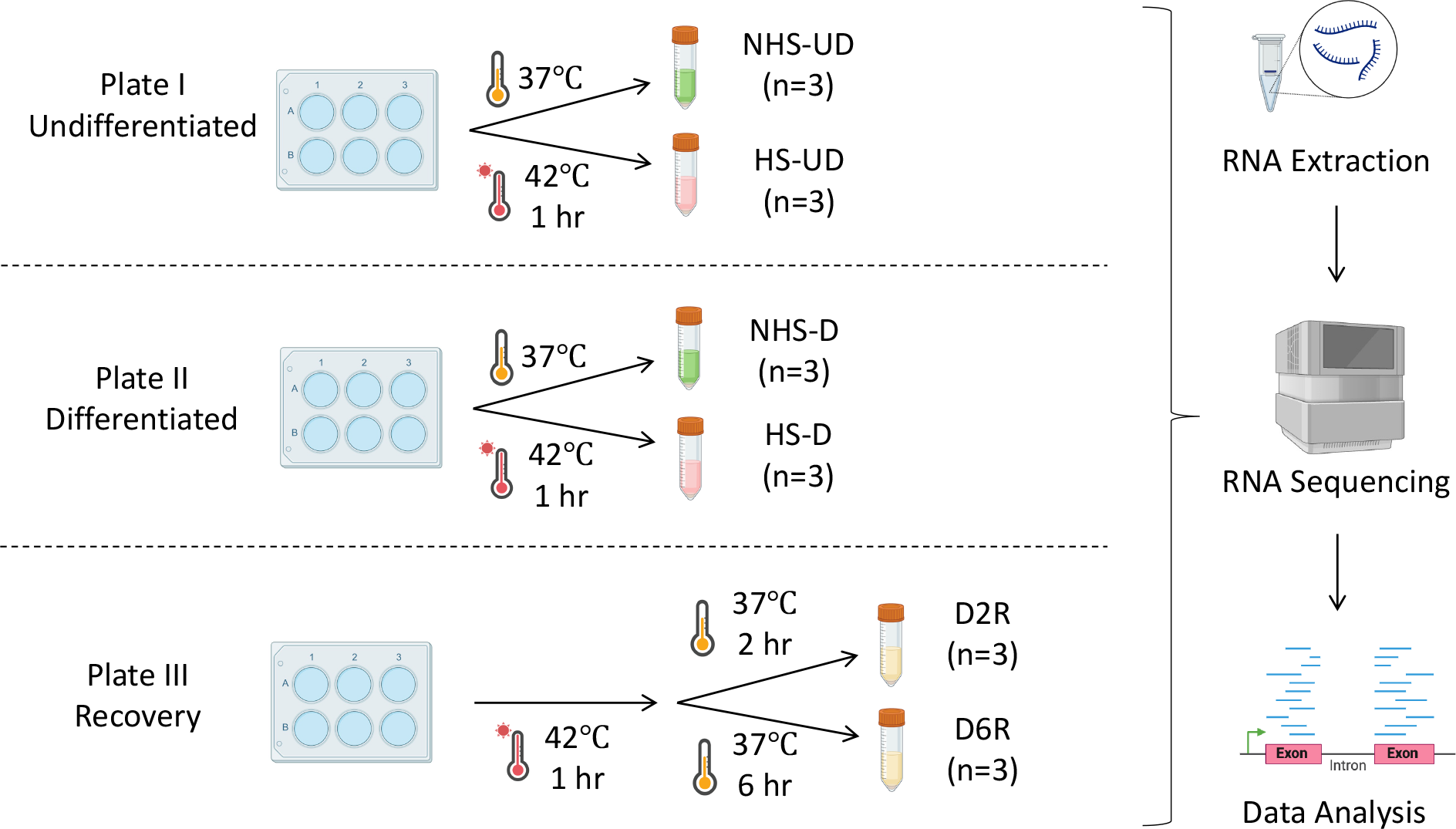
Experimental design for cell culture. All groups were cultured at 37℃ in basal or lactogenic medium for four days until collection. Plate I: Undifferentiation groups were incubated at a standard temperature of 37℃ (NHS-UD) or heat shocked at 42℃ (HS-UD) for 1 hour before collection. Plate II: Differentiated cells were incubated at a standard temperature of 37℃ (NHS- D) or heat shocked at 42℃ (HS-D) for 1 hour before collection. Plate III: Differentiated recovery cells were heat shocked for 1 hour at 42℃, then allowed to recover at 37℃ for 2 (D2R) or 6 (D6R) hours before collection. Icons were adapted from BioRender.com. NHS-UD: non-heat stress undifferentiated group; HS-UD: heat stress undifferentiated group; NHS-D: non-heat stress differentiated group; HS-D: heat stress differentiated group; D2R: heat stress differentiated 2 hour recovery group; D6R: heat stress differentiated 6 hour recovery group.

**Figure 2.**
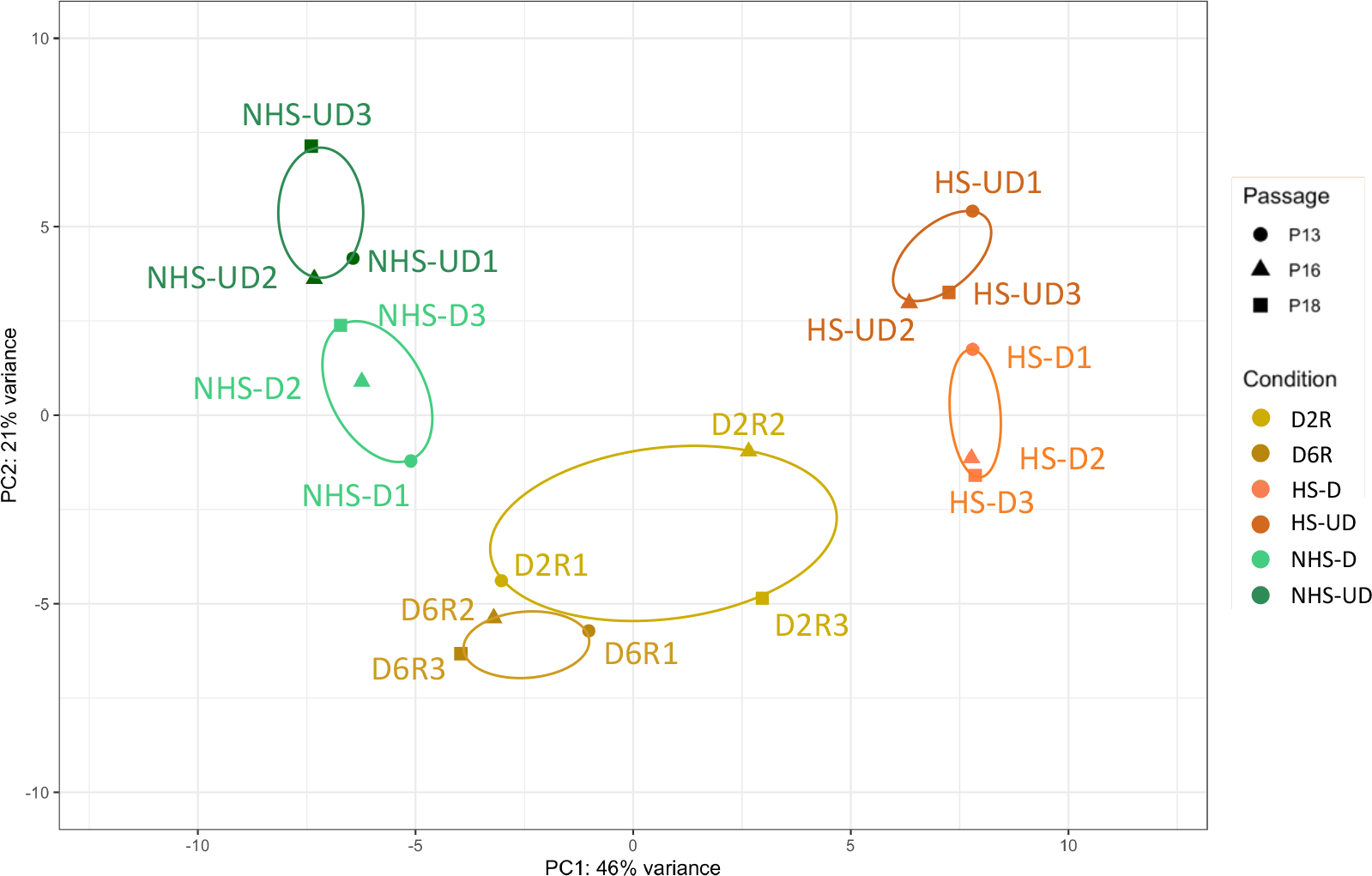
*Biological* replicate-based grouping in principal component analysis (PCA). Each point represents a sample. Circles, triangles, and squares represent cell passage 13, 16, and 18, respectively, in each repeated experiment. Colors correspond to conditions: D2R: heat stress differentiated 2 hour recovery group; D6R: heat stress differentiated 6 hour recovery; HS-D: heat stress differentiated group; HS-UD: heat stress undifferentiated group; NHS-D: non-heat stress differentiated group; NHS-UD: non-heat stress undifferentiated group.

**Table 1.** Differentially expressed genes (DEGs) in pair-wise comparisons.

| Pairwise Comparison | Total DEGs | Up-regulated DEGs | Down-regulated DEGs |
| --- | --- | --- | --- |
| NHS-UD vs. NHS-D | 154 | 92 | 62 |
| HS-UD vs. HS-D | 193 | 126 | 67 |
| NHS-UD vs. HS-UD | 349 | 156 | 193 |
| NHS-D vs. HS-D | 349 | 144 | 205 |
| HS-D vs. D2R | 151 | 106 | 45 |
| HS-D vs. D6R | 472 | 357 | 115 |

NHS: non heat stress MAC-T cells; UD: undifferentiated MAC-T cells; D: differentiated MAC-T cells; HS: heat stress exposed MAC-T cells; D2R and D6R: differentiated heat stress MAC-T cells with a 2 or 6 hour recovery, respectively.

### Transcriptomic impacts of differentiation on MAC-T Cells

A total of 154 DEGs were found in the comparison between non-heat stress (HS) undifferentiated (UD) (NHS-UD) and non-HS differentiated (D) (NHS-D) groups, showing that prolactin did affect the MAC-T cells. In the NHS-D group, we identified 92 upregulated genes, including solute carrier family 14 member 1 (*SLC14A1*), keratin 83 (*KRT83*), and tripartite motif containing 63 (*TRIM63*), and 62 downregulated genes, such as anterior gradient protein 2 homolog (*AGR2*) and neurocalcin delta (*NCALD*) (Fig. 3A, Additional file 3: Table S3). The effect of differentiation under the HS condition was also demonstrated by the comparison between HS-UD and HS-D groups, from which a total of 193 DEGs, 126 upregulated and 67 downregulated, were detected in D MAC-T cells (Fig. 3B, Additional file 3: Table S3). For example, MHC class II antigen (*BOLA-DQB*) and iodothyronine deiodinase *2* (*DIO2*) were upregulated while synaptotagmin 1 (*SYT1*) was downregulated.

**Figure 3.**
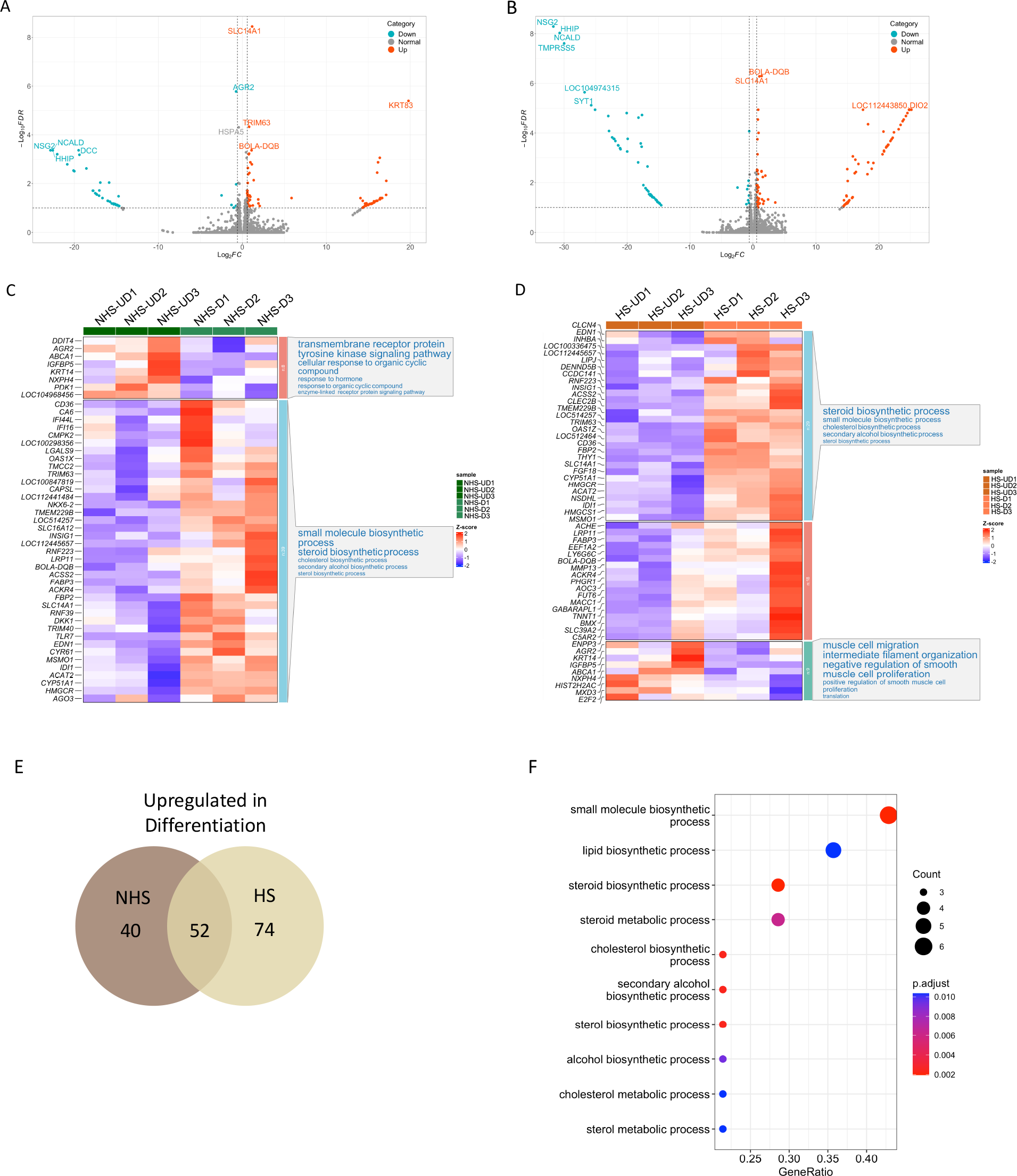
Differentiated MAC-T cells were enriched in genes and pathways related to lipid metabolism. Volcano plots of NHS-UD versus NHS-D (A) and HS-UD versus HS-D (B) comparisons representing fold change along the X-axis and false discovery rate on the y-axis; blue represents downregulated genes, grey are unchanged and red represents upregulated genes. Heatmap of relative gene expression, gene clustering, and GO enrichment analyses of top DEGs in NHS-UD versus NHS-D (C) and HS-UD versus HS-D (D). Higher expression is indicated by red, whereas lower is blue. Venn diagram of DEGs upregulated in differentiated groups (E) and corresponding GO terms (F). NHS-UD, non-heat-stressed undifferentiated group; NHS-D, non- heat-stressed differentiated group; HS-UD, heat-stressed undifferentiated group; HS-D, heat- stressed differentiated group.

Top DEGs from both comparisons were clustered and subjected to Gene Ontology (GO) analysis (Fig. 3C-D). In both NHS and HS conditions, D MAC-T cells were enriched for genes related to small molecule, steroid, and cholesterol biosynthetic processes, while undifferentiated cells were enriched for genes involved in basic cellular functions such as cell proliferation and translation. Fifty-two DEGs, including *SLC14A1*, fatty acid binding protein 3 (*FABP3*), acyl-CoA synthetase short chain family member 2 (*ACSS2*), and fatty acid translocase (*CD36*), were upregulated in D MAC-T cells in both comparisons, regardless of HS condition (Fig. 3E, Additional file 3: Table S3). These common DEGs are associated with lipid, steroid, and cholesterol biosynthetic, and metabolic processes (Fig. 3F). Differential expression of milk protein genes was not detected by RNA-seq in this study. Three milk protein genes (*CSN1S1*, *CSN2*, *LALBA*) were detected by RT-qPCR, but only *CSN2* showed a significant upregulation in differentiated cells under thermal-neutral condition (Additional file 4: Fig. S1).

### Transcriptomic impacts of heat stress (HS) on MAC-T Cells

To study the effect of HS on MAC-T cell gene expression, we first used UD cells and compared NHS-UD and HS-UD groups. In the HS group, 349 DEGs were identified, with 156 upregulated and 193 downregulated genes (Fig 4A, Additional file 5: Table S4). Several molecular chaperones, including *HSPA6*, *DNAJB1*, *HSPH1,* and *HSPA1A*, were upregulated, while genes related to epithelial cell function, fibrinogen like 2 (*FGL2*), homeobox D4 (*HOXD4*), and milk yield-associated genes, BarH like homeobox 1 (*BARHL1*) and transfer RNA cysteine (*TRNAC-ACA*), were downregulated. Next, the effect of HS on D MAC-T cells was determined by the comparison between NHS-D and HS-D groups. A total of 349 DEGs were identified with 144 upregulated genes and 205 downregulated genes in the HS group (Fig. 4B, Additional file 5: Table S4). Similar upregulation of HSPs was observed while Krüppel-like factor 10 (*KLF10*), polo like kinase 2 (*PLK2*), and solute carrier family 7 member 5 (*SLC7A5*) were downregulated. GO analysis from both comparisons (NSH-UD versus HS-UD and NHS-D versus HS-D) showed that HS cells were enriched in genes related to protein folding, chaperone-mediated protein folding, response to heat, and temperature stimulus, while genes involved in RNA metabolism and transcription regulation by RNA polymerase II were enriched under NHS conditions (Fig. 4C-D). The top 5 enriched Kyoto Encyclopedia of Genes and Genomes (KEGG) pathways (Fig. 4E-F, Additional file 6: Table S5) of HS cells were related to the cellular response to HS, including protein processing in endoplasmic reticulum (ER), legionellosis, and antigen processing and presentation. On the other hand, pathways enriched in NHS groups were associated with protein synthesis, cell differentiation, and cell cycle regulation, including mTOR, TNF, and FoxO signaling pathways.

**Figure 4.**
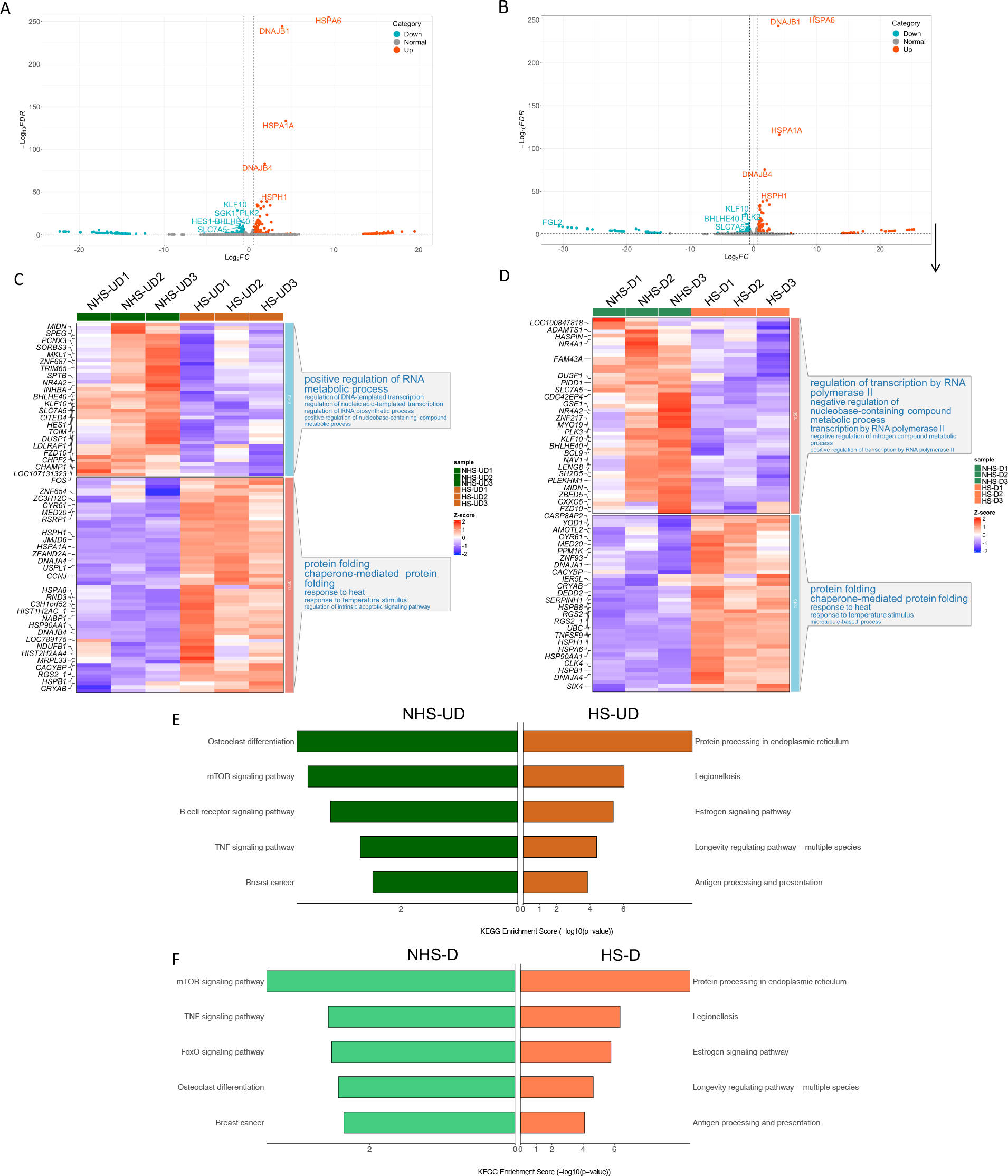
MAC-T cells showed a strong heat shock response. Volcano plots of NHS-UD versus HS-UD (A) and NHS-D versus HS-D (B) comparisons representing fold change along the X-axis and false discovery rate on the y-axis; blue represents downregulated genes, grey are unchanged and red represents upregulated genes. Heatmaps of relative gene expression, gene clustering, and GO enrichment analysis of top differentially expressed genes (DEGs) in NHS-UD versus HS-UD (C) and NHS-D versus HS-D (D). Higher expression is indicated by red, whereas lower is blue. KEGG enrichment analysis of top DEGs in NHS-UD versus HS-UD (E) and NHS-D versus HS- D (F). NHS-UD: non-heat stress undifferentiated group; NHS-D: non-heat stress differentiated group; NHS-UD: heat stress undifferentiated group; HS-D: heat stress differentiated group.

To identify the common effect of HS on MAC-T cells cultured with or without differentiation medium, we analyzed the up- or downregulated DEGs from these two comparisons (NHS-UD versus HS-UD and NHS-D versus HS-D). After HS, 96 downregulated genes and 81 upregulated genes were found to overlap (Additional file 7: Fig. S2A-B). For example, *HSP*s, Jun proto-oncogene (*JUN*), sodium/myo-inositol cotransporter (*SLC5A3*), and FKBP prolyl isomerase 4 (*FKBP4*) were activated by HS, while *FGL2, BARHL1, TRNAC-ACA, JUNB*, *SLC7A5,* and serum/glucocorticoid regulated kinase 1 (*SGK1)* were repressed. GO analysis showed the overlapping genes that were down regulated after HS were related to cell differentiation, response to endogenous stimulus and growth factors, and positive regulation of RNA metabolism. In contrast, overlapping genes upregulated after HS were associated with strong molecular function regulation, protein folding chaperone, heat shock protein binding and ATP-dependent activity (Additional file 7: Fig. S2C-D).

In addition, the enriched motif of transcription factors (TFs) that are potential direct regulators in the promotor regions of HS-downregulated and HS-upregulated genes were predicted. The top 60 TFs identified are shown in Table 2. Among the TFs upregulated by HS, two heat shock factors (HSFs), HSF1 and HSF4, were found. In addition, Krüppel-like factors (KLFs) family, including KLF5, KLF4, KLF7, KLF9, KLF12, and KLF14, were also enriched. KLF5, KLF12, KLF14, and zinc finger and BTB domain containing family 4 (ZBTB4) and 7B (ZBTB7B) were identified as enriched motifs in both HS-downregulated and HS-upregulated gene promoters.

**Table 2.** Transcription factors with motifs enriched in the promoter region of heat stress (HS)-activated and HS-repressed genes.

| **HS-activated** | **P val.** | **FDR** | **HS-repressed** | **P val.** | **FDR** |
| --- | --- | --- | --- | --- | --- |
| **KLF5** | 1.16E-05 | 0.00348533 | **E2F3** | 1.25E-07 | 6.09E-05 |
| **EGR4** | 1.43E-05 | 0.00348533 | **TFAP2D** | 2.63E-07 | 6.09E-05 |
| **KLF7** | 3.47E-05 | 0.0043483 | **E2F2** | 3.74E-07 | 6.09E-05 |
| **KLF12** | 4.37E-05 | 0.0043483 | **E2F1** | 4.99E-07 | 6.09E-05 |
| **EGR3** | 4.45E-05 | 0.0043483 | **ZNF202** | 8.63E-07 | 8.44E-05 |
| **KLF4** | 5.50E-05 | 0.00448217 | **SP2** | 1.40E-06 | 0.00010245 |
| **GLIS1** | 7.50E-05 | 0.00523839 | **PLAG1** | 1.47E-06 | 0.00010245 |
| **SP1** | 0.00010623 | 0.00649309 | **EGR1** | 3.34E-06 | 0.00020396 |
| **MLL** | 0.00027864 | 0.01382177 | **E2F4** | 4.29E-06 | 0.00023325 |
| **SP6** | 0.00028265 | 0.01382177 | **TFAP2C** | 5.65E-06 | 0.00027647 |
| **E2F3** | 0.00040089 | 0.01782159 | **ZIC4** | 9.12E-06 | 0.00038298 |
| **KLF9** | 0.00063441 | 0.02585203 | **E2F6** | 9.40E-06 | 0.00038298 |
| **ZBTB4** | 0.00079988 | 0.02795234 | **EGR3** | 1.20E-05 | 0.00045128 |
| **ZNF202** | 0.00080027 | 0.02795234 | **SP1** | 1.56E-05 | 0.00053016 |
| **GMEB2** | 0.00105852 | 0.03450784 | **SP6** | 1.63E-05 | 0.00053016 |
| **SP2** | 0.00144316 | 0.04410646 | **ZBTB4** | 2.07E-05 | 0.00063356 |
| **KDM2B** | 0.00190178 | 0.05470426 | **SP8** | 2.21E-05 | 0.00063674 |
| **SP8** | 0.00235226 | 0.06390315 | **TFCP2L1** | 2.93E-05 | 0.00079681 |
| **BRCA1** | 0.0025152 | 0.0647334 | **AP2A** | 7.82E-05 | 0.00201344 |
| **EGR1** | 0.00299421 | 0.07004097 | **KLF14** | 9.73E-05 | 0.00233024 |
| **E2F2** | 0.00300789 | 0.07004097 | **TFAP2B** | 0.00010007 | 0.00233024 |
| **HSF1** | 0.00337375 | 0.07498925 | **GLIS1** | 0.00010691 | 0.00236175 |
| **LOC514433** | 0.00385812 | 0.08202705 | **ZBTB7B** | 0.00011108 | 0.00236175 |
| **KLF14** | 0.00449214 | 0.09152743 | **NHLH1** | 0.00012558 | 0.0024654 |
| **ZBTB7B** | 0.00471318 | 0.09218979 | **GCM1** | 0.00012604 | 0.0024654 |
| **GMEB1** | 0.00562069 | 0.10571214 | **Q8SQA** | 0.00016661 | 0.0030849 |
| **TFCP2L1** | 0.00668306 | 0.11897435 | **MLL** | 0.00017215 | 0.0030849 |
| **HSF4** | 0.00681244 | 0.11897435 | **PAX5** | 0.00017664 | 0.0030849 |
| **PLAG1** | 0.00747968 | 0.1261229 | **KLF12** | 0.00018809 | 0.00317165 |
| **TFEB** | 0.00814934 | 0.13283425 | **KLF5** | 0.00019576 | 0.00319094 |

P val.: p-value; FDR: false discovery rate.

### Transcriptomic recovery responses of differentiated (D) MAC-T cells after heat stress (HS)

To investigate the potential of MAC-T cells to recover post HS, we included two recovery (R) groups of D MAC-T cells. One group was returned to culture at 37°C for 2 (D2R) and the other for 6 hours (hrs) (D6R). When comparing D2R to the HS-D group, 151 DEGs were identified, with 106 upregulated and 45 downregulated genes (Fig. 5A, Additional file 8: Table S6). For example, *JUNB*, *SGK1*, and *HSP90B1* were upregulated, while C3 and PZP like alpha-2- macroglobulin domain containing 8 (*CPAMD8*), T-box transcription factor 15 (*TBX15*), and integrin beta2 (*ITGB2*) were downregulated after 2 hr R. After 6 hrs R from HS, 472 DEGs were identified when compared with the HS-D group. We identified 357 upregulated genes, including glutamate-cysteine ligase catalytic subunit (*GCLC*), WD repeat and SOCS box containing 1 (*WSB1*), and DNA damage-inducible transcript 4 (*DDIT4*), and 115 downregulated genes, such as *HSP6A, DBAJB1,* and *HSPA1A* in the D6R group (Fig. 5B, Additional file 8: Table S6). GO terms related to cell death and apoptosis processes in cells were enriched after 2 hr recovery, and GO terms associated with gene functions in protein modification and hypoxia response were upregulated after 6 hr recovery (Fig. 5C-D).

**Figure 5.**
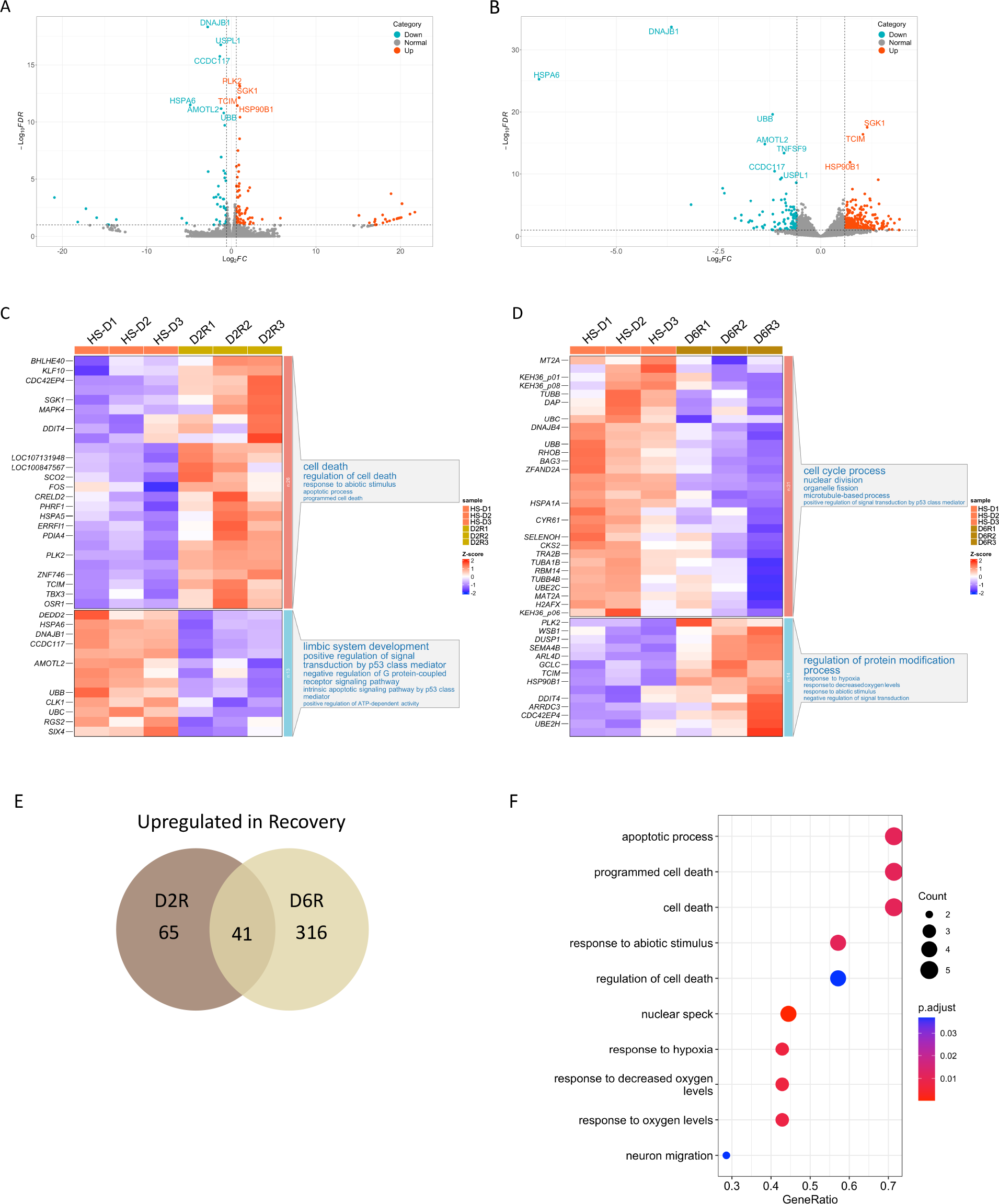
Differentiated MAC-T cells showed dynamic responses during post-HS recovery. Volcano plots of HS-D versus D2R (A) and HS-D versus D6R (B) comparisons representing fold change along the X-axis and false discovery rate on the y-axis; blue represents downregulated genes, grey represents unchanged, and red represents upregulated genes. Heatmaps of relative gene expression, gene clustering, and GO enrichment analysis of top differentially expressed genes (DEGs) in HS-D versus D2R (C) and HS-D versus D6R (D). Higher expression is indicated by red, whereas lower is blue. Venn diagram of DEGs upregulated in recovery groups (E) and corresponding GO terms, where the size of the dot represents the count and the color corresponds to the adjusted P-value. (F). HS-D: heat stress differentiated group; D2R: heat stress differentiated 2 hour recovery group; D6R: heat stress differentiated 6 hour recovery group.

When overlapping DEGs detected by these two comparisons, 18 genes were found to be upregulated in HS-D cells, including *HSP6A, DBAJB,* and *EDN1,* while 41 genes were elevated in both R groups, including ubiquitin specific peptidase 54 (*USP54*)*, HSP90B1,* and *SGK1* (Fig. 5E, Additional file 8: Table S6, Additional file 9: Fig. S3A). Genes upregulated during the R process were involved in apoptotic processes, response to abiotic stimulus, and response to hypoxia (Fig. 5F), while genes upregulated in HS condition were related to heat stress responses and cellular and organelle membrane components (Additional file 9: Fig. S3B).

### Transcriptomic changes of MAC-T cells under differentiation (D), heat stress (HS), and during recovery (R)

To generate a comprehensive profile of the transcriptomic dynamics of MAC-T cells under D and HS, as well as during R, we conducted clustering analysis using the top 30 DEGs from each pair- wise comparison, ensuring the removal of duplicates. This analysis resulted in the generation of six gene clusters, each associated with enriched GO terms based on the genes within that cluster (Fig. 6, Additional file 10: Table S7). Differentiation treatment led to the upregulation of genes involved in several metabolic and biosynthetic processes (Cluster 1). When stimulated by HS, cells showed a strong transcriptomic response to heat, temperature stimuli, and protein localization to mitochondrion (Cluster 6). During HS recovery, gene expression transitioned from response to ER stress and apoptotic process after 2hr (Cluster 3) to negative regulation of G protein-coupled receptor signaling pathway at 6hr (Cluster 4 and 5). Moreover, in non-HS groups or during R (Cluster 2), cells expressed genes enriched in the positive regulation of cell differentiation and developmental process (Fig. 6).

**Figure 6.**
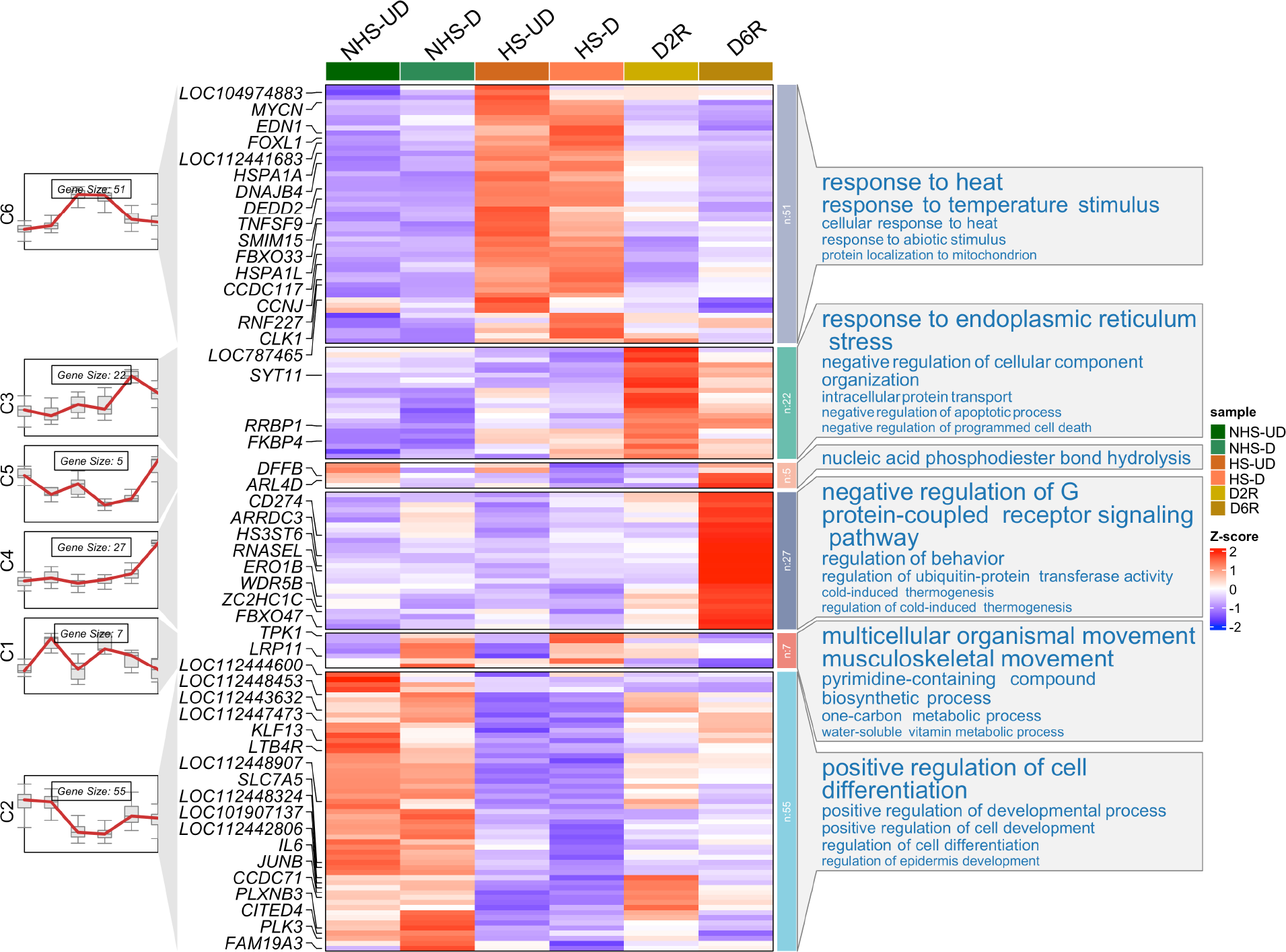
Transcriptomic dynamics of MAC-T cells under heat stress and during recovery periods. Line plots, heatmaps of relative gene expression, gene clustering, and GO enrichment analysis of top differentially expressed genes (DEGs). Higher expression is indicated by red, whereas lower is blue. NHS-UD: non-heat stress undifferentiated group; NHS-D: non-heat stress differentiated group; HS-UD: heat stress undifferentiated group; HS-D: heat stress differentiated group; D2R: heat stress differentiated 2 hour recovery group; D6R: heat stress differentiated 6 hour recovery group.

### RT-qPCR validation of RNA-seq data

Nine genes (*FBP2*, *TLR7*, *THY1*, *HSPH1*, *DNAJB1*, *JUNB*, *SGK1*, *ABCA1*, *SOX2*) that were differentially expressed in pair-wise comparisons, as identified by RNA-seq, were selected and expressions levels were analyzed by RT-qPCR (Addition file 11: Fig. S4). Fold changes of gene expression from both RNA-seq and RT-qPCR showed the same up- or down-regulation trend, suggesting that our RNA-seq data accurately captured the expression of genes from samples.

## Discussion

In this study, we generated a comprehensive profile of transcriptome dynamics of bovine MAC-T cells after differentiation, exposure to HS, and recovery (Fig. 7). We found an upregulation of genes involved in lipid metabolism in differentiated MAC-T cells, which also showed a strong response to heat stimulus under HS and exhibited the upregulation of genes involved in chaperon-mediated protein folding activities. HS repressed genes associated with cellular differentiation, transcription, and RNA metabolism. During recovery, MAC-T cells exhibited ER stresses and apoptotic process at 2 hours post HS. After 6 hours, cells shifted to express genes related to protein modification and hypoxia response.

**Figure 7.**
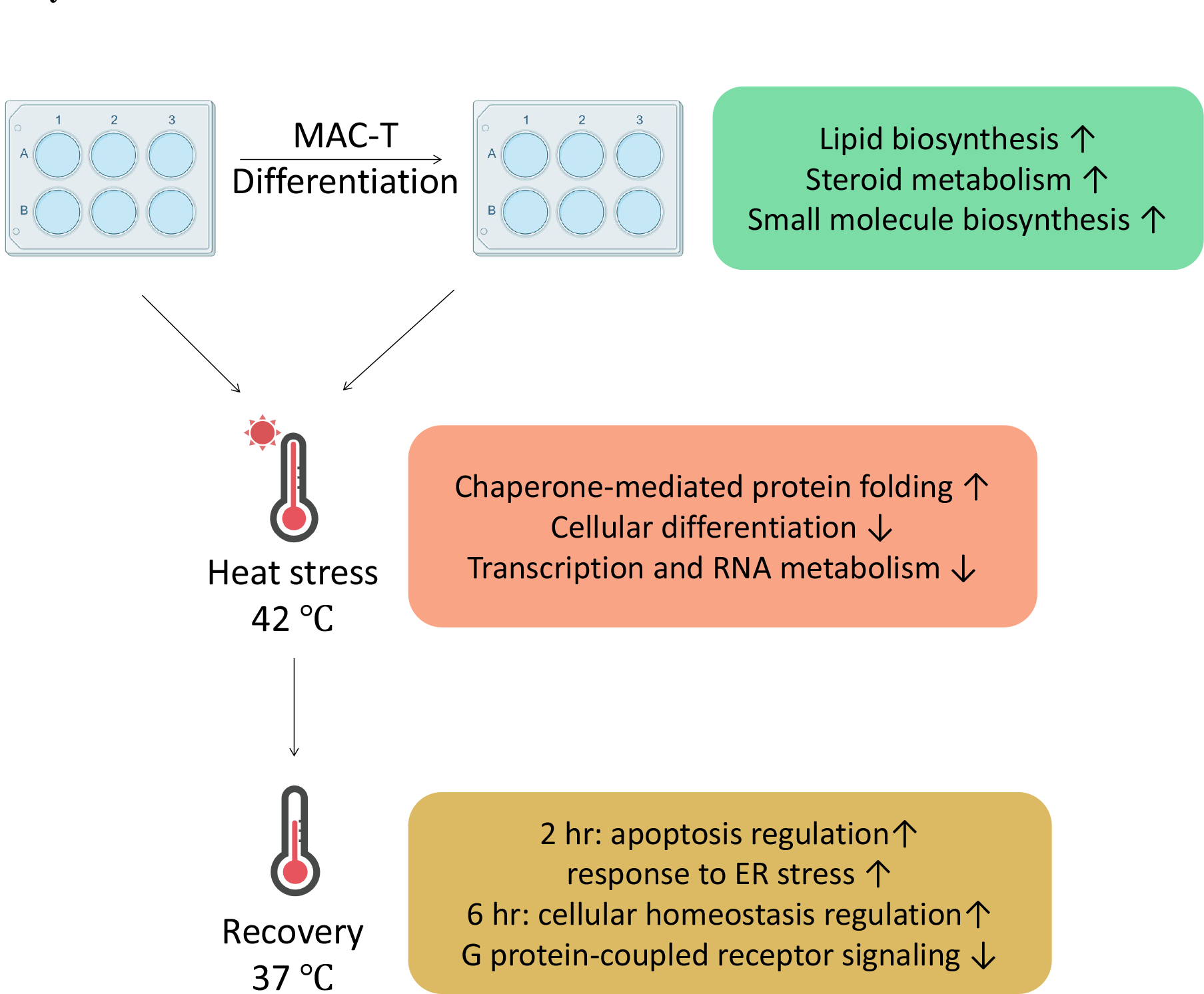
Enriched GO terms of MAC-T cells under differentiation, heat stress, and recovery. After differentiation, MAC-T cells exhibited increased expression of genes related to lipid synthesis. Heat stress triggered expression of genes related to protein folding activity while reduced expression of genes tied to differentiation and RNA metabolism. After 2 hour and 6 hour recovery, differentiated MAC-T cells had upregulated expression of genes involved in regulation of apoptosis and response to hypoxia. Icons were adapted from Biorender.com.

The differentiation of mammary epithelial cells is critical for preparing dairy cattle for lactation ^20,21^. This process, often indicated by casein (*CSN1*, *CSN2*) production, involves complex hormones *in vivo* and requires supplements in the culture medium ^22,23^. However, consistent differentiation of MAC-T remains challenging due to variability in hormones and culture conditions ^24^. Our research showed that under non-HS conditions MAC-T cells increased *CSN2* expression in a lactogenic medium as detected by qPCR (Additional file 4: Fig. S1), but not by RNA-seq. This could be attributed to the differences between these two technologies, or due to the use of high-glucose in medium (4.5 g/L), which promotes cell growth but may suppress casein and hormone receptor gene expression ^25^. However, we observed upregulation of a solute carrier family 14 member 1 (*SLC14A1*) in differentiated cells. *SLC14A1* is essential for transporting urea, which is a key milk component and the main breakdown product of protein metabolism of lactating cow mammary tissues ^26^. Additionally, several genes involved in milk fat synthesis were also upregulated, including fatty acid binding protein 3 (*FABP3*), fatty acid transferase (*CD36*), and acyl-CoA synthetase short chain family member 2 (*ACSS2*), suggesting active lipid metabolism in MAC-T response to the differentiation medium ^27^. *FABP3* and *CD36* are essential for long-chain fatty acid uptake in mammary cells, while *ACSS2* activates short-chain fatty acids, all three show high expression during lactation ^28–30^. Moreover, *FABP3* is the direct target of sterol regulatory element-binding protein (SREBP), which is a key secretory activator of lipid biosynthesis in milk production ^28^. These findings align with previous studies that support the use of MAC-T cells for investigating milk fat production pathways, especially those regulated by SREBP ^13^. Moreover, GO analysis highlighted the enrichment of biosynthetic processes that are crucial for cellular function and milk production in differentiated MAC-T cells ^31^. Thus, our data indicate a positive response of MAC-T cells to the differentiation medium, consistent with expected gene expression profiles for milk lipid synthesis.

HS triggers a cascade of cell protection mechanisms, including the transcription of heat shock proteins (HSPs) driven by heat shock factor 1 (HSF1) to maintain protein stability and cellular functions under stress ^3^. Vertebrates have four HSF members (HSF1-4), with HSF1 as the primary regulator ^32,33^. In our study, HSF1 and HSF4 were predicted to enrich in the promoter regions of HS-activated genes. Previous works showed increased *HSF1* gene expression in undifferentiated MAC-T cells ^14,16^ and dairy cattle blood cells ^34^ under HS. HSF4 is less well-defined but has found upregulated in the heart and liver of laying chickens under HS ^35^.

HSPs, molecular weights ranging from small (HSP20) to large (HSP110), play key roles in protein folding and cellular protection ^36,37^. Our study found induced HSP genes expression in HS conditions, such as *HSPB1* (HSP20), *DNAJB1* (HSP40), *HSPA6* (HSP70), *HSP90AA1* (HSP90), and *HSPH1* (HSP110). Additionally, GO and KEGG analysis showed enrichment of chaperone-mediated protein folding, protein processing in ER pathways, and estrogen signaling pathways, which associated with heat tolerance traits in dairy cattle ^38^.

In addition to HSFs and *HSP*s, DEGs altered by HS also enriched the binding sites of Krüppel-like factors (KLFs) family and Zinc finger and BTB domain containing (ZBTB) family in our study. KLF4 protein was found to regulate the continuous expression of HSP90 in murine cells ^39^. In a chicken cell line, *KLF5* gene expression was found upregulated after HS ^40^. Moreover, ZBTB7B, known as the vertebrate homolog of GAGA-associated factor (GAF) ^41,42^, was identified as a main HS regulator in gene activation but also contributes to repression in *Drosophila* ^43^. Overall, the upregulation of HSFs and other known HS regulators confirmed the HSR of differentiated MAC-T cells, and ZBTB7B could play a dual function in dairy cattle HSR regulation.

Our results show that HS similarly impacts both differentiated (D) and undifferentiated (UD) cells, disrupting protein folding and RNA metabolism and downregulating solute carrier family 7, member 5 (*SLC7A5*), an essential amino acid transporter for milk synthesis ^44,45^. Downregulation of *SLC7A5* was also observed in undifferentiated MAC-T cells ^15^ and in dairy cows ^10^ under HS. In addition, we found that HS repressed transfer RNA cysteine (*TRNAC*-*ACA*) and BarH like homeobox 1 (*BARHL1*), which are related to milk yield and fat percentage, respectively ^46,47^. Moreover, our KEGG analysis also revealed that non-HS groups enriched on mTOR pathway genes, which are essential for milk protein synthesis but reduced by HS in dairy cow MG ^10^. Overall, our results show a strong cellular response to HS that affected differentiation in MAC-T cells.

Considerable research efforts have focused on HS response in dairy cattle, yet recovery from HS and its residual effects remain less explored. Studies using mammary epithelial cells showed reduced cell viability after 3 hours and reached the lowest (∼70% of cells) after 6 hours of recovery ^48,49^. Another study observed significantly altered expression of *HSPs* and milk genes after HS treatment, which gradually returned to the normal level after 24 hours ^19^.

In our study, post-HS recovery for 2 hours showed enriched genes for anti-apoptotic processes and ER stress regulation, including *HSP90B1* and *JUNB*. *JUNB* mediates anti-apoptotic properties by repressing ER stress in pancreatic beta cells ^50^. In contrast, *JUN*, an inducer of apoptosis ^51^, was downregulated during recovery. Moreover, T-box transcription factor 15 (*TBX15*), which is related to temperature adaptation in cattle ^52^, was also downregulated after 2 hours recovery. Our result suggested that HS triggers apoptosis in MAC-T cells, and as a response, a 2 hour recovery activated genes in anti-apoptotic function and ER stress mitigation.

After 6 hours of recovery, the transcription of MAC-T shifted to cellular homeostasis regulation, such as protein modification and hypoxia responses. Oxidative markers genes, including WD repeat and SOCS box containing 1 (*WSB1*) and glutamate-cysteine ligase catalytic subunit (*GCLC*) ^53,54^, and mitochondrial function genes, including PPARG coactivator 1 alpha and beta (*PPARGC1A* and *PPARGC1B*) ^55,56^, were upregulated. This is consistent with previous observation in mice that HS caused hypoxia and oxidative stress ^57^. Therefore, our results suggest that MAC-T cells switched to cellular homeostasis regulation after 6 hours of recovery from HS.

Common genes upregulated at both 2 and 6 hours recovery included *HSP90B1*, DNA damage-inducible transcript 4 (*DDIT4*), polo like kinase 2 (*PLK2*), and serum- and glucocorticoid-inducible kinase 1 (*SGK1*). *HSP90B1* was the only HSP gene that maintained a high expression level after the 6 hour recovery, indicating its role in ER stress mediation and protein folding, consistent with previous findings in bovine ^19^ and rat ^58^ cells. *DDIT4* and *PLK2* were linked to oxidative stress and mitochondrial dysfunction ^59,60^, while *SGK1*, which is involved in several cellular processes ^61–63^, responds to HS and mitigate oxidative stress in human cells ^64^. HSP90 also enhances SGK1 activity, which is crucial in MECs survival ^65,66^. Overall, our results demonstrated that MAC-T cells undergo a dynamic recovery from HS that encompasses regulation of apoptosis and cellular homeostasis at 2 and 6 hours.

## Conclusions

Our study revealed the dynamic transcriptomic profiles of MAC-T cells under HS and during recovery. MAC-T cells showed increased gene expression for lipid metabolism after differentiation. HS induced a response in protein folding activity, impaired MAC-T cell differentiation and RNA metabolic processes. Several transcription factors were identified as potential regulators for HSR in MAC-T cells. Recovery marked by the transcriptomic shifts from apoptotic regulation to cellular homeostasis regulation. These data provided a comprehensive profile of the MAC-T transcriptome in HS and during recovery and revealed new insights into the cellular dynamics of bovine MAC-T cells under HS.

## Methods

### Cell culture, heat stress, and recovery treatments

The bovine mammary alveolar cell (MAC-T) line ^12^ was a generous gift from Dr. Yves Boisclair (Department of Animal Science, Cornell University, Ithaca, NY). MAC-T cells were cultured in basal medium consisting of Dulbecco’s Modified Eagle’s Medium (DMEM, high glucose) (Corning Inc., Corning, NY), 10% fetal bovine serum (FBS) (Corning Inc.), 1% penicillin-streptomycin (MilliporeSigma, Burlington, MA), 1% L-glutamine (Life Technologies, Carlsbad, CA), 5 μg/ml insulin (Life Technologies), and 1 μg/ml hydrocortisone (Sigma-Aldrich, St. Louis, MO). MAC-T were grown in T75 flasks at 37°C with 5% CO2. When 80% confluent, cells were transferred to three PureCoat^TM^ 6-well plates (Corning Inc.) 2 x 10^5^ cell/well and divided into six groups, with 3 wells per group serving as technical replicates (Fig. 1). Plate I was seeded with non-HS and HS undifferentiated (UD) MAC-T cells and served as the NHS-UD and HS-UD group, respectively. Plate II was seeded with non-HS and HS differentiated (D) MAC-T cells and served as the NHS-D and HS-D group, respectively. Plate III was seeded with HS D MAC-T cells that underwent either 2 or 6 hours (hrs) recovery (R), resulting in the D2R and 6DR group, respectively. This whole experiment was repeated three times using cells at passage 13, 16, and 18, respectively.

The UD groups were cultured in basal medium and the D groups were cultured in lactogenic medium, as previously described with modifications ^23^. Briefly, lactogenic medium consisted of basal medium supplemented with 5% FBS, 5 μg/ml ovine prolactin (Protein Laboratories Rehovot Ltd., Israel), and 10 μg/ml dexamethasone (Sigma-Aldrich). All groups were cultured for four days and media were changed daily. The non-HS groups were incubated at 37°C until collection and the HS groups were incubated at 42°C for 1 hr before collection on the fourth day. The R groups were cultured at 37°C for another 2 or 6 hr after HS treatment. All cells were collected by dissociation with a 0.25% trypsin/EDTA solution (Thermo Fisher Scientific, Waltham, MA). Cell pellets were stored in 1 mL TRIzol reagent (Thermo Fisher Scientific) and kept at -80°C until RNA extraction.

### RNA extraction

Total RNA was extracted from cell pellets as described in our previous study ^8^. Briefly, cell pellets were homogenized in TRIzol reagent (Thermo Fisher Scientific). Lysates were transferred to Phasemaker tubes (Thermo Fisher Scientific) with chloroform for a 20 min centrifugation at 4°C at 14,000 RPM. The aqueous phase was then washed and eluted using RNeasy Mini Kit (Qiagen, Germany) according to the manufacturer’s instructions. Concentrations of RNA samples were determined by a NanoDrop 1000 spectrophotometer (Thermo Fisher Scientific), and samples with 260/280 and 260/230 ratios of approximately 2 were considered pure. RNA quality was determined by an Agilent 2100 Bioanalyzer (Agilent Technology, Santa Clara, CA). Each sample had an RNA integrity number (RIN) greater than 9, confirming high quality. Two technical replicates with high concentration and quality were pooled from each of the six groups in three repeated experiments, resulting in a total of 18 samples submitted for RNA sequencing.

### Library construction and data analysis of RNA sequencing

The quality control, library construction, and RNA sequencing were conducted by Novogene Inc. (Sacramento, CA) on Illumina platforms to generate paired-end raw reads.

From the raw sequencing data, adaptors were removed, and quality control was conducted using fastp (v0.23.2) ^67^. Reference genome of bos taurus (ARS-UCD 1.3) was downloaded from the UCSC database, and alignment was conducted using STAR (v2.7.9a) ^68^. FeatureCounts (v2.0.3) was used to extract raw read counts of genes ^69^. Gene expression level normalization was performed using the *DEseq2* R package ^70^, and top 300 variable genes were used to conduct principal component analysis (PCA). The batch effect was removed by *limma* R package ^71^.

To identify differentially expressed genes (DEGs), the raw read count matrix was processed by *DEseq2* R package ^70^ with differentiation, HS, and cell passage information used as experiment factors. Across six groups, pair-wise comparisons were conducted and DEGs were identified. The selecting criteria of DEGs included |Fold Change (FC)| > 1.5 and adjusted p-value < 0.1. Next, k-means clustering of top DEGs, heatmaps, and gene ontology (GO) enrichment analysis were performed using *ClusterGVis* R package ^72^. Kyoto Encyclopedia of Genes and Genomes (KEGG) pathway enrichment of DEGs were conducted using clusterProfiler (v4.0.5) ^73^. Motif search for promoter TF analysis was performed using ShinyGo (v0.77) ^74^. The promoter regions were defined as 300bp upstream of transcription start site (TSS) of DEGs.

### Real-time quantitative PCR validation

Real-time quantitative polymerase chain reaction (RT-qPCR) was performed to validate the results of RNA-seq. Three milk protein genes (casein alpha subunit 1*, CSN1S1;* casein beta*, CSN2;* lactalbumin alpha*, LALBA*) were detected. Nine genes (fructose-bisphosphatase 2*, FBP2;* Toll-like receptor 7, *TLR7*; Thy-1 cell surface antigen*, THY1;* heat shock protein family H (Hsp110) member 1*, HSPH1*; DnaJ Heat Shock Protein Family (Hsp40) Member B1*, DNAJB1*; Jun B proto-oncogene*, JUNB*; glucocorticoid-regulated kinase 1*, SGK1;* ATP-binding cassette transporter*, ABCA1;* SRY-Box transcription factor 2*, SOX2*) with differential expression larger than 1-fold in RNA-seq were randomly selected. iScript cDNA Synthesis Kit (Bio-Rad, Hercules, CA) was used to reverse transcribe one *μ*g total RNA according to the manufacturer’s instructions. SsoAdvanced Universal SYBR Green Supermix (Bio-Rad) was used to conduct RT-qPCR using two technical replicates in a CFX384 Touch Real-Time PCR machine (Bio-Rad) according to the manufacturer’s instructions. The primers were designed using NCBI Primer-BLAST (Additional file 1: Table S1). The internal control gene was glyceraldehyde-3-phosphate dehydrogenase (*GAPDH*) and the relative expression was calculated with the 2^-ΔΔCT^ method ^75,76^.

## List of abbreviations

NHS-UD: Non-heat stress undifferentiated group
NHS-D: Non-heat stress differentiated group
HS-UD: Heat stress undifferentiated group
HS-D: Heat stress differentiated group
D2R: Heat stress differentiated 2 hour recovery group
D6R: Heat stress differentiated 6 hour recovery group
PCA: Principal component analysis
RNA-seq: RNA sequencing
DEG: Differentially expressed gene
GO: Gene ontology
KEGG: Kyoto Encyclopedia of Genes and Genomes
FC: Fold change
GAPDH: Glyceraldehyde-3-phosphate dehydrogenase
RT-qPCR: Real-time quantitative PCR
TF: Transcription factor
HSP: Heat shock protein
HSF: Heat shock factor
FDR: False positive rate

## Declarations

### Ethics approval and consent to participate

Not applicable.

### Consent for publication

Not applicable.

### Availability of data and material

The RNA-seq raw data have been deposited at NCBI Gene Expression Omnibus database (RNA- seq: GSE249950). The current study did not generate new code, all codes used for analysis in this study can be found according to corresponding reference.

### Competing interests

The authors declare that they have no competing interests.

### Funding

This study was funded by the National Institute of Food and Agriculture, U.S. Department of Agriculture through a Hatch Grant #7003372 to JED. A grant from the Foundation for Food and Agricultural Research (FFAR) # CA20-SS-0000000004 to GVdW, which includes matching funds from Elanco Animal Health Incorporated and New York Farm Viability Institute (NYFVI).

### Authors’ contributions

Conceptualization: XY and JED; Experimental work: XY, RMH, and ND; Experimental supervision: JED and GVdW; Data analysis and interpretation: XY, GL, YF and JED; Funding: JED and GVdW; Original draft writing: XY; Revision of original draft: XY, RMH, ND, GVdW, and JED. All authors read and approved the final version of the manuscript.

## Acknowledgements

We thank Dr. Boisclair and Dr. Giesy for providing the MAC-T cell, technical assistance and discussion in this study. We thank Dr. Heather Huson and Dr. Vimal Selvaraj for providing us access to their lab equipment and facilities.

## Supplemental information titles and legends

**Additional file 1: Table S1. Gene names, accession numbers, primer pairs, and product sizes used for RT-qPCR.** *FBP2*: Fructose-bisphosphatase 2; *TLR7*: Toll-like receptor 7; *THY1*: Thy-1 cell surface antigen; *HSPH1*: Heat shock protein family H (Hsp110) member 1; *DNAJB1*: DnaJ Heat Shock Protein Family (Hsp40) Member B1; *JUNB*: Jun B proto-oncogene; *SGK1*: glucocorticoid-regulated kinase 1; *ABCA1*: ATP-binding cassette transporter; *SOX2*: SRY-Box Transcription Factor 2; *GAPDH*: glyceraldehyde-3-phosphate dehydrogenase.

Additional file 2: Table S2. RNA-sequencing data quality control.

**Additional file 3: Table S3. Genes and DEGs expressed in differentiation treatment.** NHS- UD, non-heat-stressed undifferentiated group; NHS-D, non-heat-stressed differentiated group; HS-UD, heat-stressed undifferentiated group; HS-D, heat-stressed differentiated group

**Additional file 4: Figure S1. RT-qPCR validation of MAC-T cell differentiation**. A: expression of *CSN1S1*, *CSN2*, and *LALBA* of MAC-T cells with or without differentiation. ** p <

0.05. Error bars were mean ± S.E.M. *CSN1S1*, casein alpha subunit 1; *CSN2*, casein beta; *LALBA*, lactalbumin alpha; NHS-UD, non-heat-stressed undifferentiated group; NHS-D, non-heat-stressed differentiated group.

**Additional file 5: Table S4. Genes and DEGs expressed in heat stress treatment.** NHS-UD: non-heat stress undifferentiated group; NHS-D: non-heat stress differentiated group; HS-UD: heat stress undifferentiated group; HS-D: heat stress differentiated group.

**Additional file 6: Table S5. KEGG analysis of non-HS and HS groups.** NHS-UD: non-heat stress undifferentiated group; NHS-D: non-heat stress differentiated group; HS-UD: heat stress undifferentiated group; HS-D: heat stress differentiated group.

**Additional file 7: Figure S2. Common DEGs and GO terms in non-HS and HS groups.** Venn diagram of DEGs downregulated (A) and upregulated (C) in HS groups and corresponding GO terms (B, D). NHS-UD, non-heat-stressed undifferentiated group; NHS-D, non-heat-stressed differentiated group; HS-UD, heat-stressed undifferentiated group; HS-D, heat-stressed differentiated group.

**Additional file 8: Table S6. Genes and DEGs expressed in recovery treatment.** HS-D: heat stress differentiated group; D2R: heat stress differentiated 2 hour recovery group; D6R: heat stress differentiated 6 hour recovery group.

**Additional file 9: Figure S3. Common DEGs and GO terms downregulated in recovery groups.** Venn diagram of DEGs downregulated in recovery groups (A) and corresponding GO terms (B). HS-D: heat stress differentiated group.

**Additional file 10: Table S7. DEG list for MAC-T transcriptomic dynamic analysis.** Values were normalized Z-score. NHS-UD: non-heat stress undifferentiated group; NHS-D: non-heat stress differentiated group; HS-UD: heat stress undifferentiated group; HS-D: heat stress differentiated group; D2R: heat stress differentiated 2 hour recovery group; D6R: heat stress differentiated 6 hour recovery group.

**Additional file 11: Figure S4. RT-qPCR validation of RNA-seq results.** X axis shows the validated genes, and Y axis shows relative expression log2FoldChange. The error bar stands for standard error of the mean. NHS-UD: non-heat stress undifferentiated group; NHS-D: non-heat stress differentiated group; HS-UD: heat stress undifferentiated group; HS-D: heat stress differentiated group; D2R: heat stress differentiated 2 hour recovery group; D6R: heat stress differentiated 6 hour recovery group.

## Notes

### Competing Interest Statement

The authors have declared no competing interest.

